# Photosynthetic resistance and resilience under drought and rewatering in maize plants

**DOI:** 10.1101/2020.08.03.234542

**Authors:** Miao Qi, Xiaodi Liu, Yibo Li, He Song, Feng Zhang, Zhenzhu Xu, Guangsheng Zhou

## Abstract

Abnormally altered precipitation patterns induced by climate change have profound global effects on crop production. However, the plant functional responses to various precipitation regimes remain unclear. Here, greenhouse and field experiments were conducted to determine how maize plant functional traits respond to drought, flooding, and rewatering. Drought and flooding hampered photosynthetic capacity, particularly when severe and/or prolonged. Most photosynthetic traits recovered after rewatering, with few compensatory responses. Rewatering often elicited high photosynthetic resilience in plants exposed to severe drought at the end of plant development, with the response strongly depending on the drought severity/duration and plant growth stage. The associations of chlorophyll concentrations with photosynthetically functional activities were stronger during post-tasselling than pre-tasselling, implying an involvement of leaf age/senescence in responses to episodic drought and subsequent rewatering. Coordinated changes in chlorophyll content, gas exchange, fluorescence parameters (PSII quantum efficiency and photochemical/non-photochemical radiative energy dissipation) possibly contributed to the enhanced drought resistance and resilience and suggested a possible regulative trade-off. These findings provide fundamental insights into how plants regulate their functional traits to deal with sporadic alterations in precipitation. Breeding and management of plants with high resistance and resilience traits could help crop production under future climate change.

## Introduction

Global climate change is now leading to an enhanced frequency and intensity of drought events (Dai, 2012; Trenberth *et al.*, 2014; Donat *et al.*, 2016; Diffenbaugh *et al.*, 2017), that are now placing staple crop production and food security at risk (Lobell *et al.*, 2014; Myers *et al.*, 2017; Leakey *et al.*, 2019; Kimm *et al.*, 2020). These changes, coupled with the acceleration of industrialisation and the rapid development of social economy, are now placing agricultural water resources in tighter supply across the globe. Water availability has now become a bottleneck for food production and even social and economic development, and lack of water has triggered a series of environmental and ecological problems that now threaten sustainable development of crop production and exacerbate global undernutrition (e.g., Daryanto *et al.*, 2016; Myers *et al.*, 2017; Rosa *et al.*, 2020).

Drought is one of the most crucial environmental factors constraining crop plant productivity due to its deleterious effects on leaf photosynthetic capacity, plant growth and crop productivity at regional and global scales (Lobell *et al.*, 2014; Daryanto *et al.*, 2016; Myers *et al.*, 2017). Plants that experience drought stress have their water balance destroyed and this leads to plant growth inhibition, stomatal closure, and decreases in the photosynthetic rate (e.g., Chaves *et al.*, 2003, 2009; Xu and Zhou, 2009; Gupta *et al.*, 2020). However, plants can invoke a number of regulative strategies to deal with water deficit, including extending the root system, increasing leaf thickness, and activating an antioxidative defence system (e.g., Trapeznikov *et al.*, 2003), increased leaf thickness (Sack and Grubb, 2002), provoked antioxidative defense system (Foyer and Noctor, 2005). The responses to water deficit depend on the duration, severity and time of occurrence of the drought. For example, plants may not be affected, even favoured under mild or moderate drought, but it can be limited and even damaged by severe drought (e.g., Fereres and Soriano, 2007; Xu *et al.*, 2014). Under a mild or moderate water deficit, an increased water use efficiency (WUE), improved nutritional content, and stable grain yield often can be observed, which can improve sustainable development by allowing deficit irrigation and water-saving agricultural practices (Fereres and Soriano, 2007; Geerts and Raes, 2009; Du *et al.*, 2015; Silveira *et al.*, 2020). Further exploration of crop responses to various water conditions can therefore provide critical information for optimising crop management practices, particularly under future climate change (Lobell *et al.*, 2014; Leakey *et al.*, 2019; Kimm *et al.*, 2020).

Plants exposed to drought will frequently show a restoration of their normal physiological functions when rewatered, and to a certain degree, they can compensate for the damage caused by drought by accelerating their growth and enhancing their photosynthetic capacity (e.g., Xu *et al.*, 2009, 2010; Hofer *et al.*, 2017). An antecedent condition, such as soil water availability, may also drive the post-stress responses to other abiotic factors, indicating important complexities in plant responses to environmental factors (Xu *et al.*, 2009; Guo and Ogle, 2019). This ability to regain a normal original state after being disturbed is termed resilience (Holling, 1973; Müller *et al.*, 2016. Resilience Alliance, 2020), and can be represented by the interference level, recovery time or recovery speed (Müller *et al.*, 2016; Bhaskar *et al.*, 2018; Harrison *et al.*, 2018; Resilience Alliance, 2020).

A recent report showed that a watering treatment following a drought can lead to a greater recovery of some key functional traits in plants (Harrison *et al.*, 2018). For example, both full and partial recoveries of leaf pigment and nitrogen contents were observed in drought-stressed maize plants following rewatering (Sun *et al.*, 2018). Similarly, Voronin *et al.* (2019) documented the physiological responses of maize plants to drought and rehydration. However, information is lacking regarding the changes in photosynthetic capacity and their associations with plant growth during drought and subsequent recovery upon rewatering. The increased frequency of drought due to global climate change emphasises the importance of understanding the mechanism underlying the plant responses to drought and rewatering for both theoretical and practical applications (e.g., Hofer *et al.*, 2017; Abid *et al.*, 2018; Guo and Ogle, 2019).

Drought has been an important factor in the growth of maize, the most widely grown crop in the world. Water deficit causes unstable and low yields in many maize production areas in the world, seriously hampering plant growth and causing 25–30% reductions in grain yield in some vulnerable regions (Sharp *et al.*, 2003; Ben-Ari *et al.*, 2016; Beyene *et al.*, 2016; Li *et al.*, 2019; Kimm *et al.*, 2020). For instance, the U.S. Corn Belt, the world’s biggest maize production region, is recognised as being prone to drought and is therefore sensitive to climate change (Kimm *et al.*, 2020). Similarly, the Corn Belt of Northeastern China (CBNC) is one of the major maize production regions in China and it too shows strong sensitivity to climate variations. Drought is a particularly critical factor constraining maize production in the CBNC (e.g., Liu *et al.*, 2012; Li and Sun, 2016).

Future climate change scenarios envisage an increase in the occurrences of both drought and flooding during the growth period in maize-growing regions (Roudier *et al.*, 2016; Kimm *et al.*, 2020). Thus, elucidating the maize plant responses to drought, rewatering and flooding is crucial for the development of technology for monitoring, evaluating and minimising the damage caused by drought and flood disasters. This knowledge can also provide insight to the factors that enhance resilience in maize plants, while also serving as a feasible reference for corn yield forecasting and field water management during the growing period.

The aim of the present study was to conduct greenhouse and field experiments to determine maize plant functional responses to drought, rewatering and flooding. The greenhouse experiments involved examination of these responses following different water treatments, including pre-drought, drought, rewatering and flooding. The field experiment was conducted in a large-sized rain shelter designed to grow maize plants under 4 irrigation regimes, including pre-drought and subsequent re-irrigation. Our focus was specifically on assessing the resilience of photosynthetic capacity in response to drought and rewatering. Three hypotheses were tested: i) drought and flooding can constrain photosynthetic capacity in maize plants, particularly under severe, prolonged water stress; 2) rewatering can lead to a full recovery of photosynthetic capacity with a compensatory mechanism; 3) the resilience of photosynthetic capacity depends on the degree of drought stress and the plant development stage. The findings may improve current knowledge and strengthen future quests to produce high-yield, drought-resistant and resilient crops (see also Gupta *et al.*, 2020).

## Materials and Methods

### Greenhouse experiment design

The first experimental site was located in a greenhouse (39°48’N, 116°28’E, 67 m a.s.l.), Institute of Botany, Chinese Academy of Sciences, Beijing, China. The soil was collected from field soil (0-30 cm soil profile) at Gucheng Ecological Environment and Agro-meteorology Test Station (39°08’N, 115°40’E, 15.2m a.s.l.), Baoding city, Hebei province, North China. Plastic pots (diameter 21 cm, height 25 cm) was used. The maize cultivars is Zhengdan 958, which is currently planted extensively in North China. The seeds were sown on June 28, 2017. We filled 5.5 kg of soil per pot; and each pot was applied as 2.54 g of diammonium phosphate compound fertilizer (i.e., 750 kg ha^−1^). The three seeds were sown in each of the pot with a depth of 2.5 cm. Soon afterwards, only one healthy plant was left before the third leaf of seedlings emerged. The seedlings grew in the greenhouse with a day/night mean temperature of *c.* 28.0/20.0 °C and maximum photosynthetic photon flux density (*PPFD*) of 1,000 μmol m^−2^s^−1^).

The greenhouse experiment used four water treatments: 1) Control treatment: the soil relative water content (SRWC) was maintained at 65–75% throughout the whole experimental period. 2) Persistent drought stress: SRWC was reduced beginning at the three-leaf stage and extending to jointing stages to the SRWC of the permanent wilting point (PWP). 3) Flooding treatment: waterlogging stress was induced at the three-leaf stage and extended until the jointing stage. 4) Drought-rehydration treatment: SRWC was reduced initially at the three-leaf stage to 35% of SRWC (the leaves wilted and the lowermost leaves began to turn yellow and withered); the plants were then rewatered to 65–75% of SRWC.

The field experimental site was located at the Jinzhou Ecology and Agricultural Meteorology Center, Liaoning, Northeastern China (N 41°49’, E 121°12’, 27.4m a.s.l.). The mean annual temperature and the mean annual precipitation over 40 years were 9.9 °C and 564 mm, respectively, with an average monthly temperature of 20.9 °C and a total precipitation of 468 mm during plant growing season. The soil is characterised as medium loam type, pH 6.3, with 1.8% organic matter and a soil bulk density of 1.61 g·cm^−3^ at the 0–100 cm soil profile. The field capacity (FC) and PWP were 22.3% and 6.5% (v/v), respectively. The soil had an organic carbon content of 10.44 g kg^−1^, total nitrogen content of 0.69 g kg^−1^, phosphorus content of 0.50 g kg^−1^, and potassium content of 22.62 g kg^−1^ The planting date and maturity date were late April and late September, respectively (Song *et al.*, 2018; Li *et al.*, 2019).

The field experimental design was as detailed previously (Li *et al.*, 2019). In brief, an electric-powered waterproof shelter (4 m in height) set up in the maize field was used to establish the various precipitation regimes that we desired (details in Li *et al.*, 2019). In total, 15 plots (15 m^2^, 5 m length, 3 m width) were under the rain shelter when it rained. The following three irrigation regimes were set up: a control (i.e. the normal irrigation every 7 day); moderate drought (water withholding for 20 days); and severe drought (water withholding for 27 days from the tasselling to milking stages). In this design, irrigation water was supplied at 296, 246, and 221 mm across the maize plant growing period. The SRWC at 0-50 cm soil depth was monitored to reach severe drought levels at a range of 30–40% at the end of rainfall-withholding, whereas under normal irrigation, the SWRC was maintained at levels of 70–80% in the control and rewatering plots. The maize cultivar was Danyu 39, with a seed sowing rate of 6.0 plants m^−2^ to ensure a planting density of 4.5 plants m^−2^. A compound fertiliser (accounting for 28%, 11%, and 12% of N, P_2_O_5_, and K_2_O, respectively), applied at a *c.* 750 kg·ha^−1^, was added before sowing (Song *et al.*, 2018; Li *et al.*, 2019).

### Leaf chlorophyll content

We examined leaf chlorophyll concentrations with a SPAD 502 chlorophyll meter (Minolta Co. Ltd, Japan). This is a nondestructive technique that provides feasible and rapid assessment of leaf relative chlorophyll concentrations (a SPAD reading value) by determining leaf transmittance of red (650 nm) and infrared (940 nm) radiation. Measurements were made on 1 July (V13, 62 days after sowing [DAS]), 12 July (VT, tasselling, 73 DAS), 20 July (R1, silking, 81 DAS), 5 August (R2, blistering, 97 DAS), 9 August (R3, milking, 101 DAS), 2 September (R4, dough, 125 DAS).

### Leaf gas exchanges and chlorophyll a fluorescence

In the greenhouse experiment, the leaf gas exchange and chlorophyll *a* fluorescence were measured with an open gas exchange system (LI-6400, LI-COR Inc., Lincoln, NE) equipped with a LI-6400-40 fluorometer. Leaves was illuminated with a red-blue LED light source. The parameters were initially obtained with acquisition software. Leaves were acclimated in the chamber for at least 15 min at 28–30 °C with a CO_2_ concentration of 400 μmol mol^−1^ and a PPFD of 1200 μmol m^−2^ s^−1^. Chlorophyll *a* fluorescence was determined with the LI-6400-40 fluorometer. The steady-state fluorescence (*F_s_*) was recorded at 1200 μmol m^−2^ s^−1^, and a second saturating pulse at ~8000 μmol photons m^−2^s^−1^ was then given to obtain the maximal light-adapted fluorescence yield 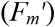. The actinic light was turned off, and the minimal fluorescence at the light-adapted state 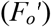 was obtained following a 3 s burst of far-red illumination.

In the field experiment, the leaf chlorophyll *a* fluorescence was determined with a miniaturised pulse-amplitude-modulated photosynthesis yield analyser (Mini-PAM, Walz, Effeltrich, Germany) equipped with a leaf clip holder (2030-Br). Light intensities (380–710 nm) were measured with the Mini-PAM microquantum sensor. After a 30 min dark adaptation, the minimal fluorescence yield (*F_o_*) was determined with a modulated light at a sufficiently low intensity (< 0.1 μmol photon m^−2^ s^−1^) to induce the minimal fluorescence. The maximal fluorescence yield (*F_m_*) was made with a 0.8 s saturating pulse at a ~7000 μmol photons m^−2^ s^−1^. The difference between the measured values (*F_m_*, *F_o_*) is the variable fluorescence (*F*_v_). Leaves were continuously illuminated at 300 μmol photons m^−1^ s^−1^ with an actinic light for 15 min. The *F_s_* was recorded, and the second saturating pulse at ~7000 μmol photons m^−2^s^−1^ was then exposed to obtain 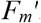. The actinic light was turned off and 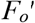 was obtained following a 3 s far-red illumination. The fluorescence parameters were calculated with the following formulas (Schreiber *et al.*, 1994; Maxwell and Johnson, 2000; Kramer *et al.*, 2004):

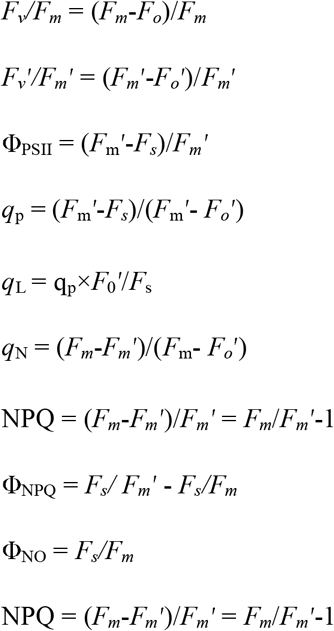

where *F_v_F_m_* is the maximal quantum efficiency of photosystem II (PSII), and 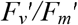 is the efficiency of excitation captured by open PSII centres in the light-adapted leaves. Φ_PSII_ is the yield of PSII photochemistry, and *q*_p_ and *q*_L_ are photochemical quenching based on puddle and lake models, respectively. NPQ or *q*_N_ is non-photochemical quenching, and both Φ_NPQ_ and Φ_NO_ are light-induced regulated non-photochemical quenching and quantum yield of non-regulated energy loss in PSII, respectively (Kramer *et al*., 2004).

### Soil relative water content

Soil was placed in an experimental pot with holes at the bottom and weighed 48 h after excessive watering to reach a saturated weight (SW) point. The soil was then dried at 110°C for at least 72 h to a constant weight (DW). The FC can be expressed as FC = (SW–DW) / DW ×100. The SRWC = Current soil water content / FC × 100.

### Resistance, recovery and resilience

Resistance was expressed as the difference/ratio in functional parameters between drought stress and ample watering as control treatments. Recovery was indicated by the difference/ratio in functional parameters between drought/pre-drought and rewatering. Resilience was calculated as the difference/ratio in functional parameters between ample watering (control treatment) and rewatering (Van Ruijven and Berendse, 2010; Ruppert *et al.*, 2015; Bhaskar *et al.*, 2018). Here, we prefer to use the percentage ratios to express these changes (Ruppert *et al.*, 2015).

### Data statistics

The data were statistically analysed with statistical software package SPSS 20.0 (SPSS Inc., Chicago, Illinois, USA). A one-way analysis of variance (ANOVA) with Duncan’s multiple comparison was used to test the differences of the functional traits between the watering treatments. The effects of watering treatment and plant/leaf developmental stages, and their interaction on the functional traits of plants, were tested with a two-way ANOVA. The correlations among the functional traits were tested with Pearson’s correlation analysis, and the relationships of photosynthetically functional traits with leaf relative chlorophyll content (SPAD readings) at different plant growth stages were tested by linear regression analysis. The comprehensive relationships between leaf photosynthetic functional traits, plant growth and their responses to irrigation regimes and plant/leaf developmental stages were determined by principal component analysis (PCA). The significance levels were set at *P* < 0.05 and 0.01, unless otherwise stated.

## Results

### Photosynthetic traits in the greenhouse

The greenhouse experiment showed that drought stress led to a slight reduction in the relative chlorophyll content (SPAD readings) 4 days after withholding water (i.e., 22 days after sowing, DAS), followed by a rise 8 days after the drought-stressed treatment (Fig. 1). However, the chlorophyll content showed a sharp linear decrease from 26 DAS to 37 DAS when the relative soil water content (RSWC) fell sharply to the severe drought stress level of 35%. After rewatering, the chlorophyll content sharply increased, with recovery values of 14.2, 15.2 and 25.6% under consecutive drought at 32, 34 and 37 DAS, respectively, indicating that a greater recovery may occur at the end of the measurement period. The resilience values were −4.3, −14.0 and −5.0% at 32, 34 and 37 DAS, respectively, showing that the resilience rose initially following rewatering, then decreased, and then increased again. Flooding led to sharp declines in SPAD after 4 days of flooding treatment, indicating that chlorophyll degradation occurred under the flooded condition.

**Fig. 1.**
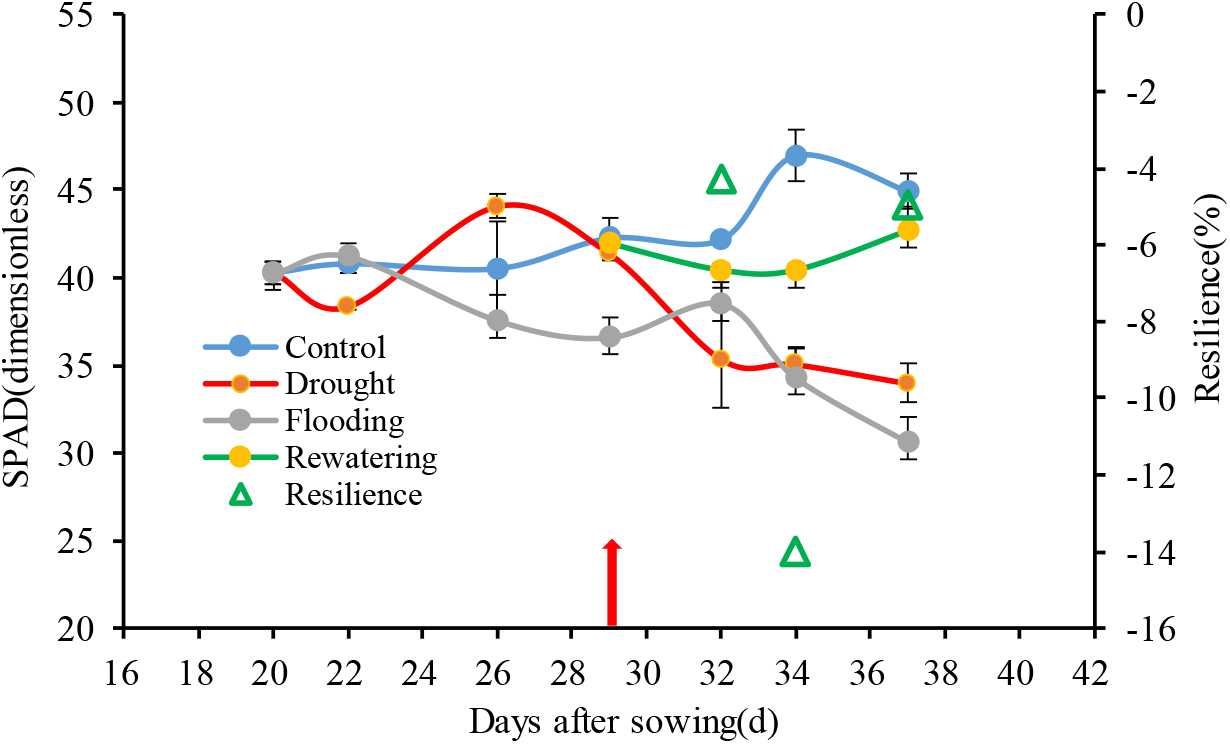
The changes in leaf relative chlorophyll content (SPAD values) in the youngest and fully expanded leaves of maize plants subjected to various watering regimes (blue line, ample watering as the control; grey line, flooding; red line, drought; green line, rewatering; green open triangle, resilience). The red arrow indicates the rewatering date; the data are shown as means ±SE (n = 3-6).

As shown in Fig. 2a, drought only led to a slight reduction in the net light-saturated photosynthetic rate (*A*_sat_) within one week after withholding water. However, this rate sharply decreased from 28.8 μmol m^−2^s^−1^ to 4.8 μmol m^−2^s^−1^ by 85.2% 29 DAS when RSWC dropped to 35%. After rewatering, *A*_sat_ sharply increased, with recovery values of 5.53, 1.18 and 5.98 times the values seen under consecutive drought at 32, 34 and 37 DAS, respectively. The rate approached and even exceeded the control level at 32, 34 and 37 DAS. The resilience values increased gradually from −12.1 to 10.2 and 25.4%, indicating a possible escalation of resilience with time after rewatering. A stimulation of the *A*_sat_ occurred during the initial 6 days under flooding; thereafter, flooding induced a decrease compared with the control treatment. However, *A*_sat_ under flooding ultimately reached the level of the control treatment. Compared with the control, stomatal conductance (*g*_s_) was significantly decreased (−96.5%) at 29 DAS, just before rewatering (Fig. 2b). A greater recovery was observed, but only positive resilience was detected at 34 DAS.

**Fig. 2.**
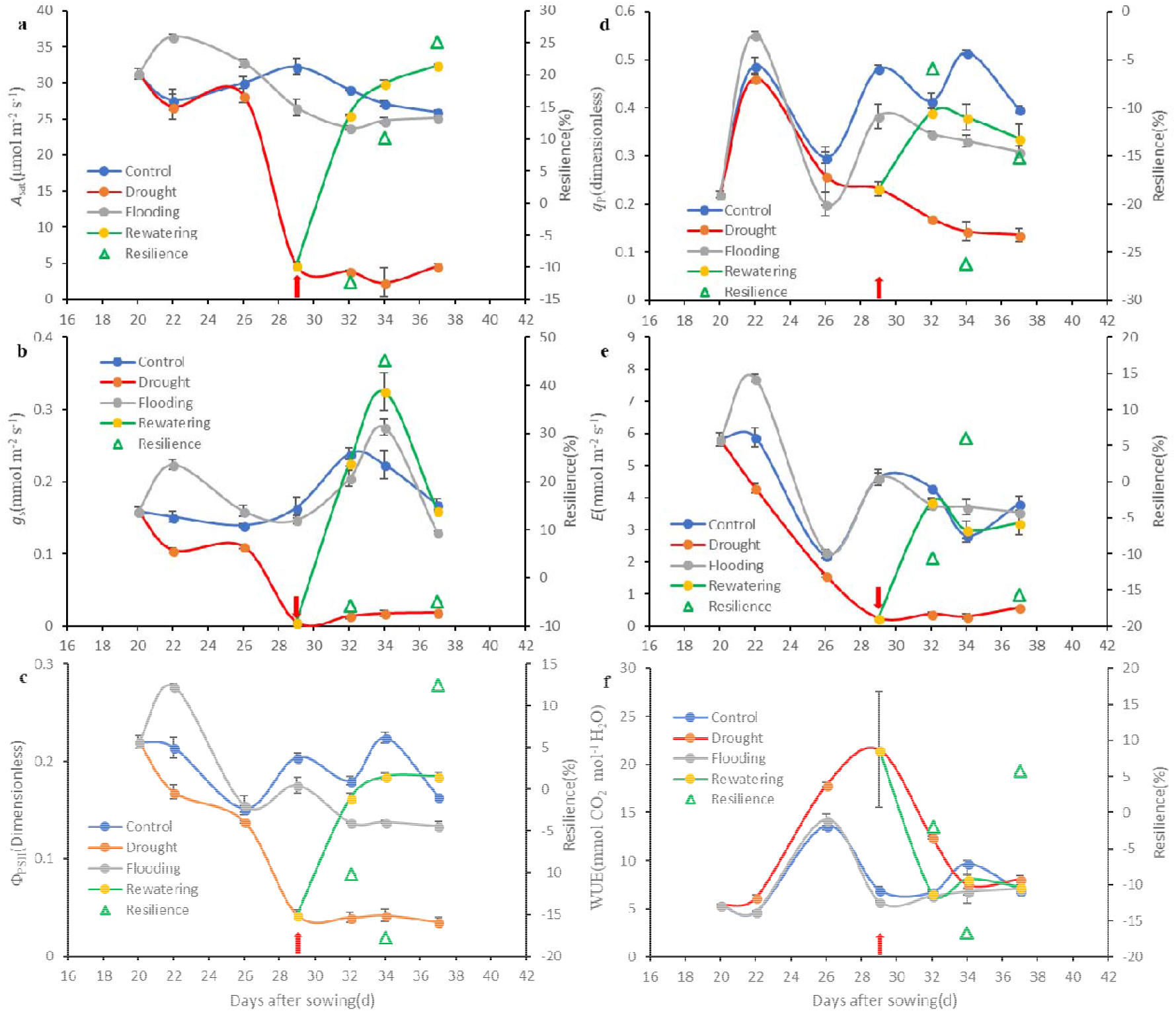
The changes in net light-saturated photosynthetic rate (*A*_sat_, a), stomatal conductance (*g*_s_, b), quantum yield of PSII electron transport (Φ_PSII_, c), photochemical quenching (*q*_P_, d), transpiration rate (*E*, e), and water use efficiency (WUE, f) in the youngest and fully expanded leaves of maize plants subjected to various watering regimes (blue line, ample watering as the control; grey line, flooding; red line, drought; green line, rewatering; green open triangle, resilience). Red arrows indicate the rewatering date; the data are shown as means ±SE (n = 3-6).

The quantum yield of PSII electron transport (Φ_PSII_) decreased with drought-treatment time, dropping to its lowest value (by 79.3%) at 29 DAS (Fig. 2c). The recovery values were 3.05, 3.38 and 4.22 times relative to continuous drought at 3, 5 and 8 days after rewatering, respectively. Flooding also led to an initial stimulation in Φ_PSII_; thereafter, the value decreased below the control level. The photochemical quenching (*q*_P_) showed a substantial fluctuation even under the control treatment (Fig. 2d). However, a dramatic decline of 51.9% was observed after 9 days of water withholding. We also found recoveries of 1.3-, 16.6- and 14.8-fold at 3, 5 and 8 days following rewatering, respectively. However, the increases still did not reach the level of the control, so the resilience values were negative (−5.74, −26.22 and −15.22 at 3, 5 and 8 days following rewatering). A stimulation of *q*_P_ was also observed initially at 2 days after flooding exposure, but this disappeared thereafter and the value dropped to levels lower than the control levels.

The transpiration rate (*E*) significantly decreased due to drought stress, dropping to the lowest point at 29 DAS (a decrease of 94.5% relative to control, Fig. 2e). Rapid decreases occurred following the rewatering, with recovery values of 9.3-, 8.9- and 4.6-fold the values under continuous drought at 3, 5 and 8 days following rewatering, respectively. However, the resilience values were −10.4, 6.0, and −15.7% at 3, 5 and 8 days following rewatering, respectively. A stimulation of *E* also appeared initially by flooding; thereafter, however, the similar *E* changes were similar to those of the control (Fig. 2e). Leaf water use efficiency (WUE) was initially increased by drought, but subsequently decreased with drought-exposure time, indicating that the enhancement of WUE may be attenuated by the water deficit intensity and its persistent duration. Rewatering led to a decline in WUE at the earlier stage, but thereafter WUE remained stable relative to both the control and continuous drought plants. WUE was not affected significantly by flooding during the experimental periods (Fig. 2f).

Similar responses were observed in the mature leaves (Fig. 3a-f). Drought reduced *A*_sat_, with great recovery and a positive resilience noted at the end of the experiment (Fig. 3a). A sharp rise appeared during the initial flooding, but *A*_sat_ decreased thereafter. A drastic *g*_s_ resilience was evident at 34 DAS (Fig. 3b). A great recovery occurred for Φ_PSII_ and *q*_p_; however, the negative resilience was still maintained (Fig. 3c, d). A rapid and drastic reduction in *E* was observed by imposition of drought stress, with great recovery; however, the resilience remained negative (Fig. 3e). Drought always elevated the WUE in the mature leaves, whereas flooding did not substantially affect it. Only a small positive resilience was observed at the end of the experiment (Fig. 3f).

**Fig. 3.**
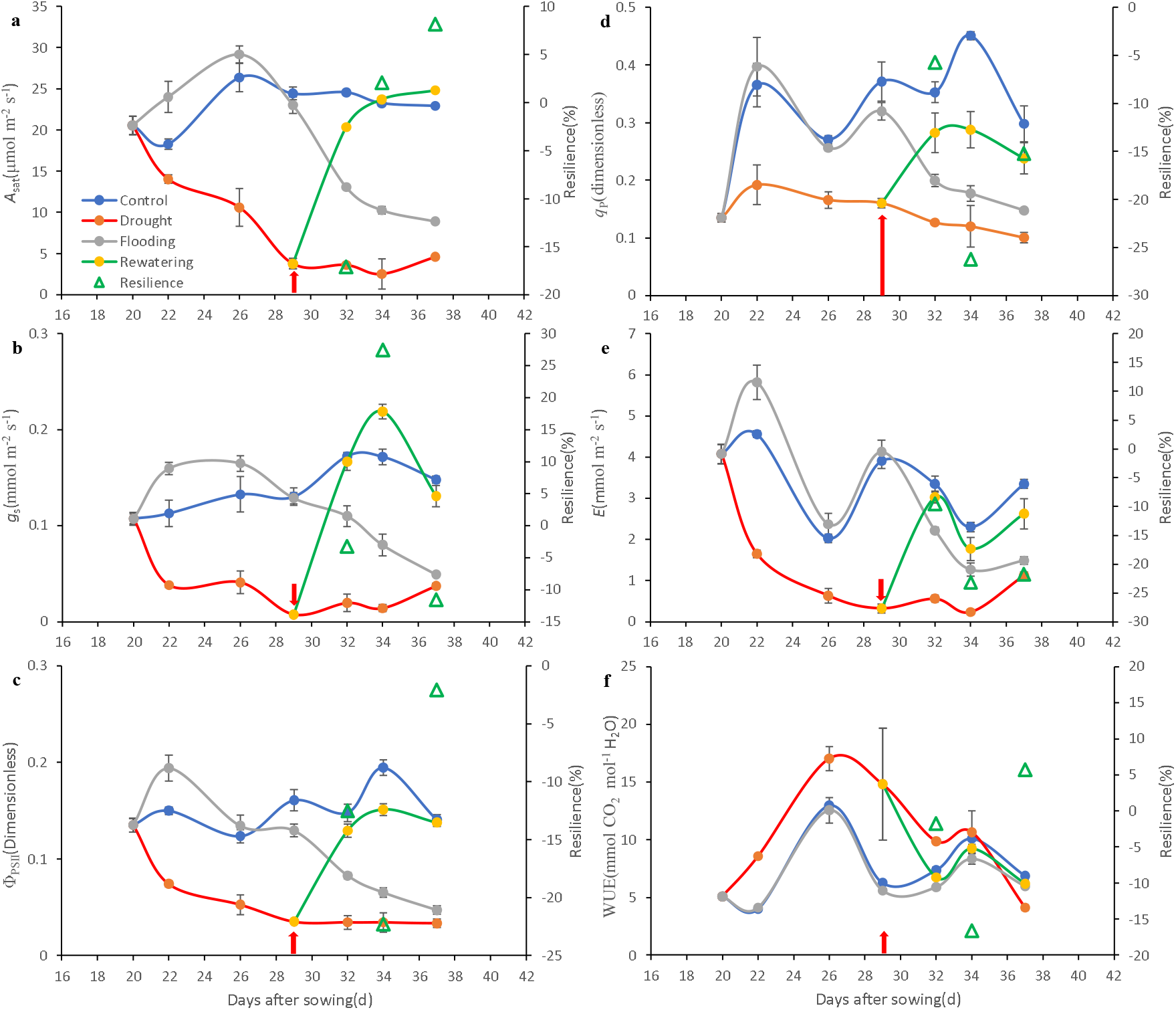
The changes in net light-saturated photosynthetic rate (*A*_sat_, a), stomatal conductance (*g*_s_, b), quantum yield of PSII electron transport (Φ_PSII_, c), photochemical quenching (*q*_P_, d), transpiration rate (*E*, e), and water use efficiency (WUE, f) in the mature leaves of maize plants subjected to various watering regimes (blue line, ample watering as the control; grey line, flooding; red line, drought; green line, rewatering; green open triangle, resilience). Red arrows indicate the rewatering dates; the data are shown as means ±SE (n = 3-6).

### Photosynthetic traits in the field

In the field experiment, the upper canopy leaves in the control treatment showed gradual increases in the relative chlorophyll content (SPAD values) from 1 July (62 DAS, V13), 12 July (73 DAS, VT, tasselling), July 20 (81 DAS, R1, silking), to August 5 (97 DAS, R2, blistering), until reaching a maximum on 97 DAS; the relative chlorophyll content then decreased as plant development progressed (Fig. 4a). Episodic droughts led to dramatic declines, whereas rewatering led to more increases (i.e., a positive recovery) under moderate drought (MD) than under severe drought (SD). Negative resilience values were observed under both drought treatments at 101 and 125 DAS. The maximum quantum efficiency of PSII (*F_v_/F_m_*) showed a similar pattern to that seen for the relationship with DAS; i.e., a unimodal curve (Fig. 4b). A drastic decline occurred under SD; however, recovery was greater following rewatering.

**Fig. 4.**
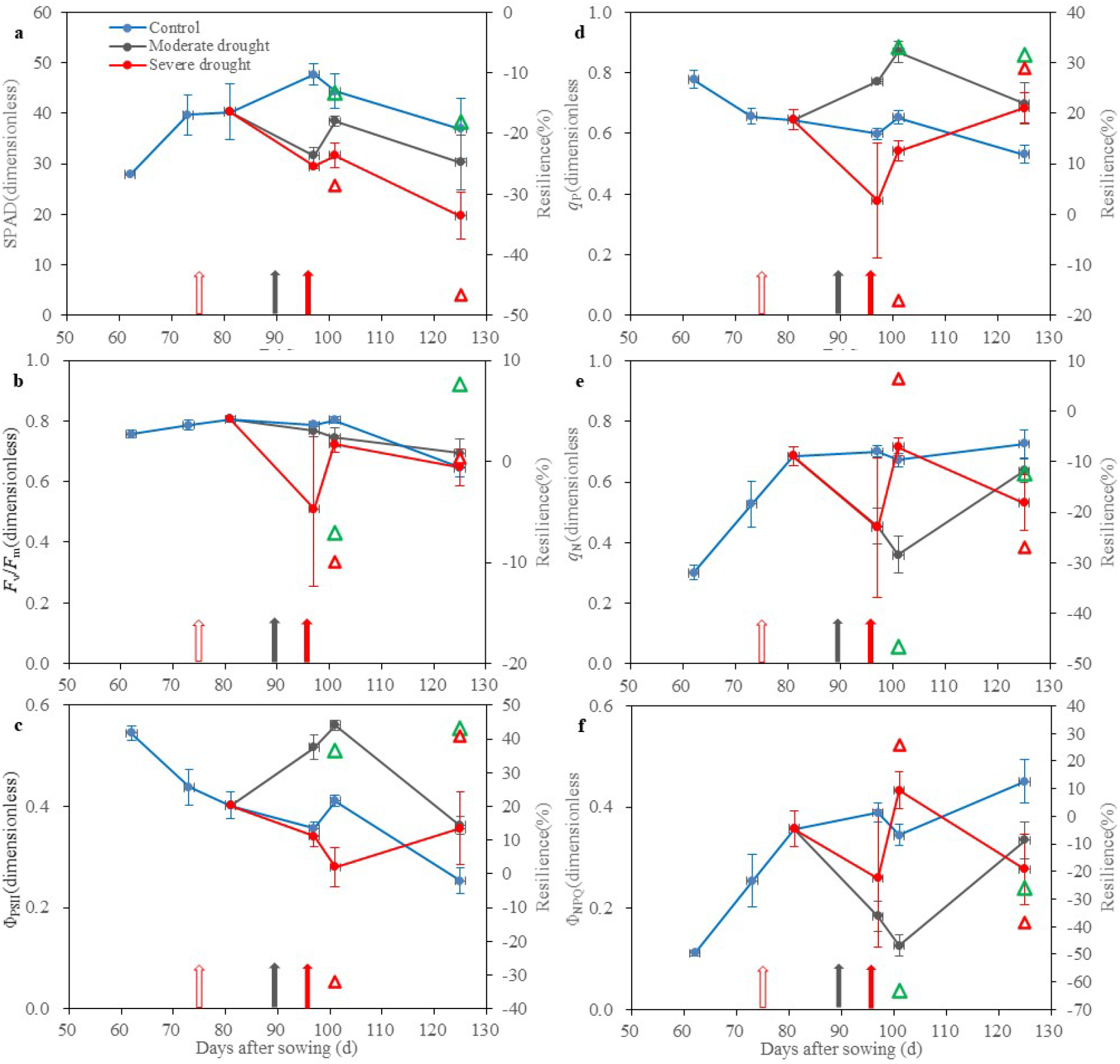
Chlorophyll *a* fluorescence in ***upper*** leaves under drought and rewatering. Green and red open triangles represent the resilience of moderate and severe drought at 101 and 125 days after sowing (DAS), respectively. Red open arrows indicate the DAS of water withholding; while grey and red close arrows indicate the rewatering DASs of moderate and severe drought treatments, respectively. The data are shown as means ±SE (n = 3-6). *F_v_/F_m_,* maximal quantum efficiency of photosystem II (PSII); Φ_PSII_, the yield of PSII photochemistry; *q*_p_, photochemical quenching based on puddle model; *q*_N_, non-photochemical quenching; Φ_NPQ_, light-induced regulated non-photochemical quenching.

A high resilience was found with MD at 125 DAS. The Φ_PSII_ values decreased with plant growth (Fig. 4b). An increase occurred under MD, but SD led to a marked decline with greater resilience at both 101 and 125 DAS. The changes in *q*_p_ and its resilience were similar to those of Φ_PSII_ (Fig. 4d). The changes in non-photochemical quenching *q*_N_) and the yield of light-induced regulated non-photochemical quenching (Φ_NPQ_) and their resilience showed the same changing trends (Fig. 4d): they increased with DAS, and MD led to a drastic decline with a high resilience at 101 DAS.

In the middle canopy leaves, SPAD values decreased under SD, and no positive resilience was observed (Fig. 5a). Positive resilience was noted for *F_v_F_m_* at 125 DAS (Fig. 5b). The Φ_PII_ and *q*_p_ values decreased with plant development under the control treatment, but greater resilience appeared under MD at 101 and 125 DAS and under SD at 125 DAS (Fig. 5c, d). Both *q*_N_ and Φ_NPQ_ increased with DAS, with greater resilience under MD at both 101 and 125 DAS (Fig. 5e, f).

**Fig. 5.**
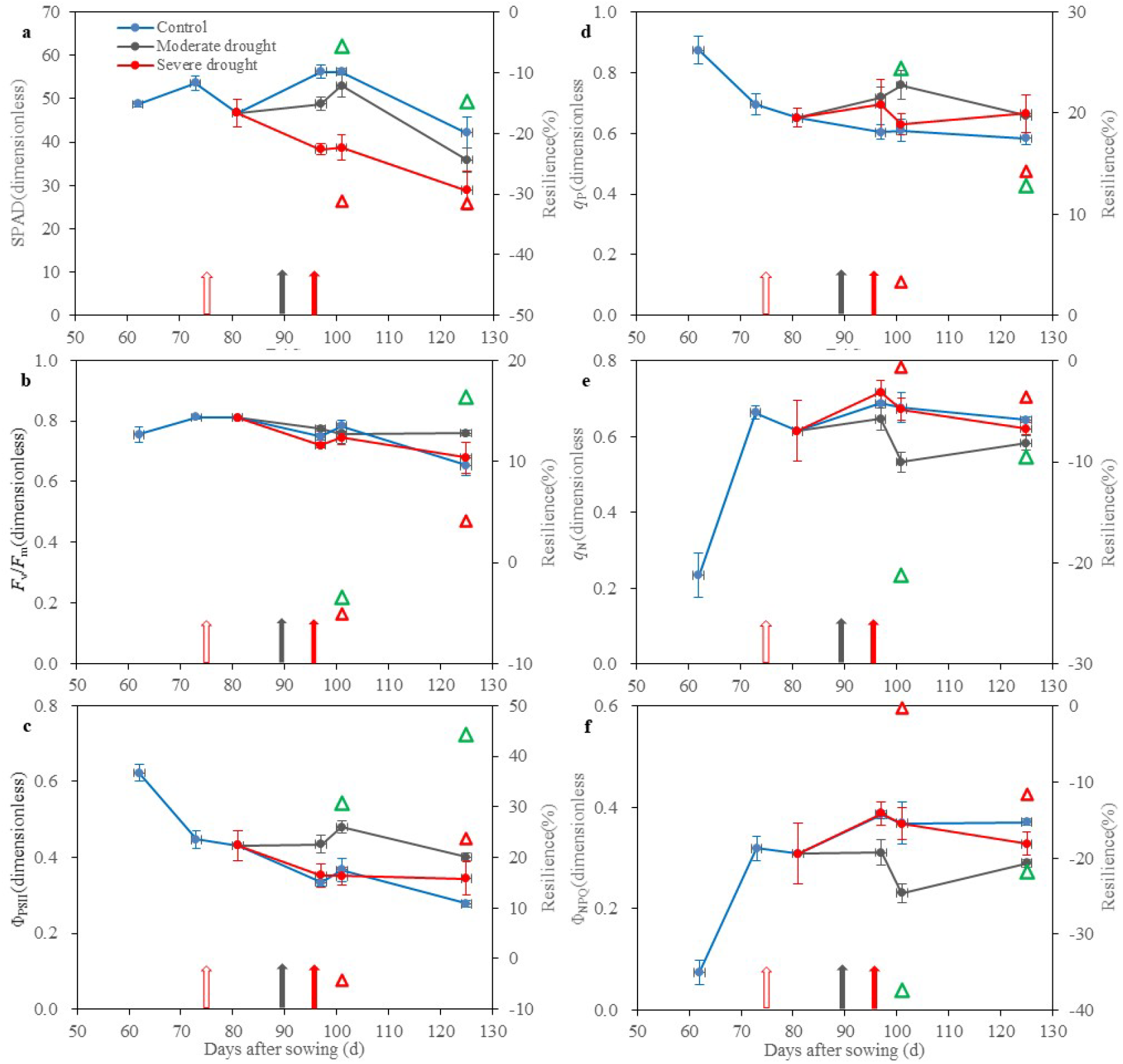
Chlorophyll *a* fluorescence in ***middle*** leaves under drought and rewatering. Green and red open triangles represent the resilience of moderate and severe drought at 101 and 125 days after sowing (DAS), respectively. Red open arrows indicate the DAS of water withholding; while grey and red close arrows indicate the rewatering DASs of moderate and severe drought treatments, respectively. The data are shown as means ±SE (n = 3-6). *F_v_/F_m_*, maximal quantum efficiency of photosystem II (PSII); Φ_PSII_, the yield of PSII photochemistry; *q*_p_, photochemical quenching based on puddle model; *q*_N_, non-photochemical quenching; Φ_NPQ_, light-induced regulated non-photochemical quenching.

In the bottom canopy leaves, the relative chlorophyll content steeply decreased with DAS under all irrigation regimes, with a marked decline under SD. Rapid recoveries occurred with rewatering; however, only negative resilience was observed (Fig. 6a). A severe drought episode resulted in a reduction in *F_v_F_m_* at 97 DAS, but a rapid recovery occurred at 4 d after rewatering (Fig. 6b). Rewatering resulted in high *F_v_/F_m_* resilience in the plants exposed to previous MD and SD at the end of plant development. A decline in Φ_PSII_ was observed from 62 to 81 DAS, but a stable Φ_PSII_ change remained thereafter during the later plant developmental periods. Marked resilience appeared for both pre-drought treatments at the two final developmental stages (Fig. 6c). The changes in *q*_p_ showed a similar pattern to that of Φ_PSII_. However, the marked resilience appeared only at 101 DAS (Fig. 6d). Under ample irrigation, both *q*_N_ and Φ_NPQ_ increased until 81 DAS and then remained stable. Drought led to declines in *q*_N_ and Φ_NPQ_ with considerable recovery at 125 DAS in the plants exposed to the SD episode; however, the resilience still remained negative (Fig. 6e, f).

**Fig.6.**
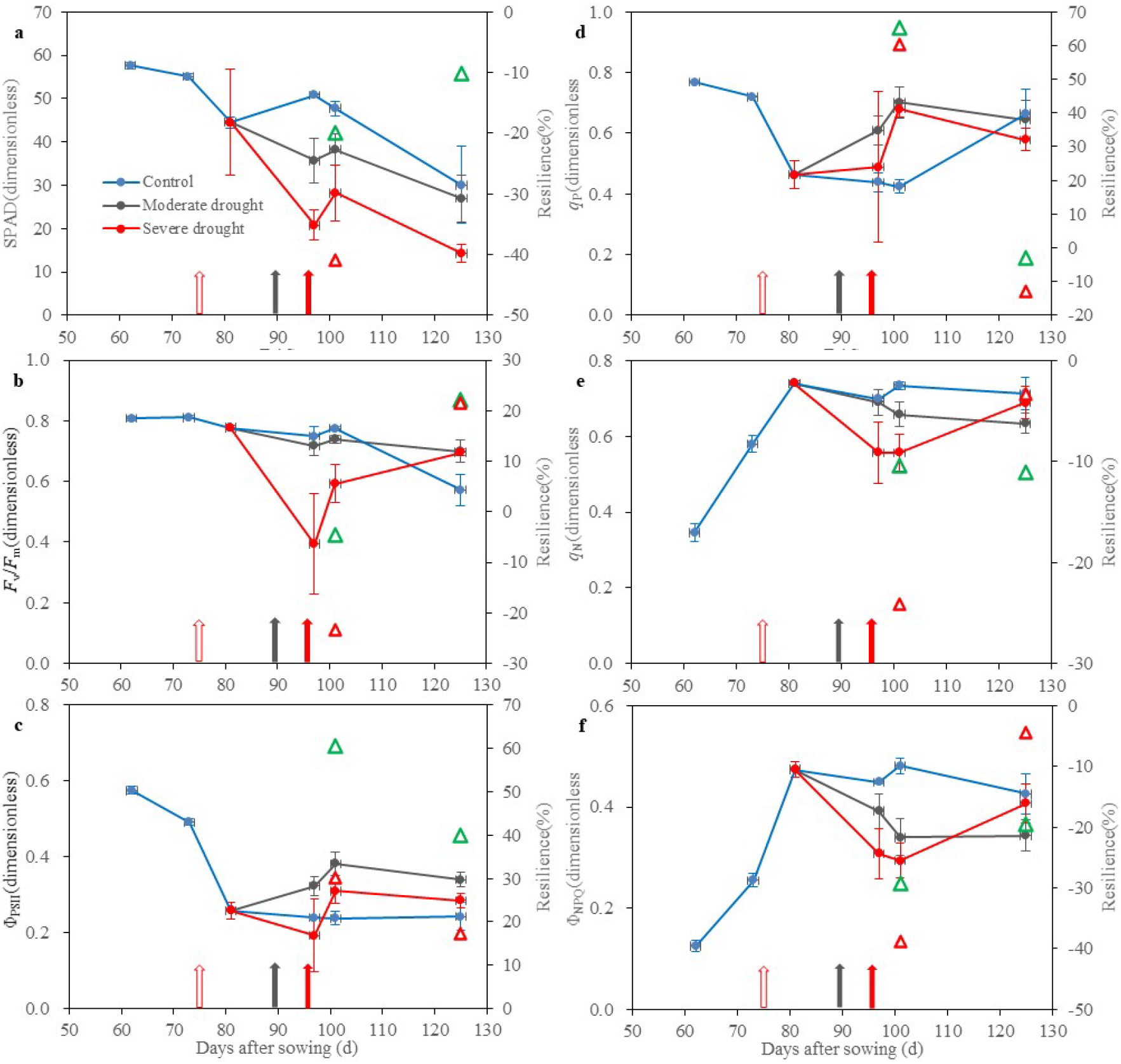
Chlorophyll *a* fluorescence in ***bottom*** leaves under drought and rewatering. Green and red open triangles represent the resilience of moderate and severe drought at 101 and 125 days after sowing (DAS), respectively. Red open arrows indicate the DAS of water withholding; while grey and red close arrows indicate the rewatering DASs of moderate and severe drought treatments, respectively. The data are shown as means ±SE (n = 3-6). *F_v_/F_m_,* maximal quantum efficiency of photosystem II (PSII); Φ_PSII_, the yield of PSII photochemistry; *q*_p_, photochemical quenching based on puddle model; *q*_N_, non-photochemical quenching; Φ_NPQ_, light-induced regulated non-photochemical quenching.

### Relationships between the functional traits

The relationships between fluorescence parameters and chlorophyll contents (SPAD values) in the maize field are shown in Fig. 7. We separated the data into sub-categories to explore how their relationships are altered at the two developmental stages. We only considered the data before/at previous tasselling stages (i.e., VT, a transitional stage from the vegetative stage to reproductive stage); therefore, the only significant and strong relationship was observed between *F_v_F_m_* and chlorophyll content (SPAD readings, R^2^ = 0.318, *P* < 0.001; Fig. 4a). The other parameters (i.e., Φ_PSII_, *q*_P_, *q*_N_ and Φ_NPQ_) showed no significant relationships (*P* > 0.05, Fig. 4c-f), except for a significant and negative relationship between *F*_s_ and the SPAD values (Fig. 4b). Using the data after VT revealed significant and positive relationships of SPAD values with fluorescence parameters, including *F*_v_*F*_m_ (R^2^ = 0.607, *P* < 0.001, Fig. 4a), *F*_s_ (R^2^ = 0.193, *P* = 0.022, Fig. 4b), Φ_PSII_ (R^2^ = 0.210, *P* = 0.016, Fig. 4c), *q*_P_ (R^2^ = 1.48, *P* = 0.047, Fig. 4d), *q*_N_ (R^2^ = 0.378, *P* = 0.001, Fig. 4e) and Φ_NPQ_ (R^2^ = 0.248, *P* = 0.008, Fig. 4f). This indicates that stronger and closer associations emerged between chlorophyll content and the key fluorescence parameters, but only at later developmental stages.

**Fig. 7.**
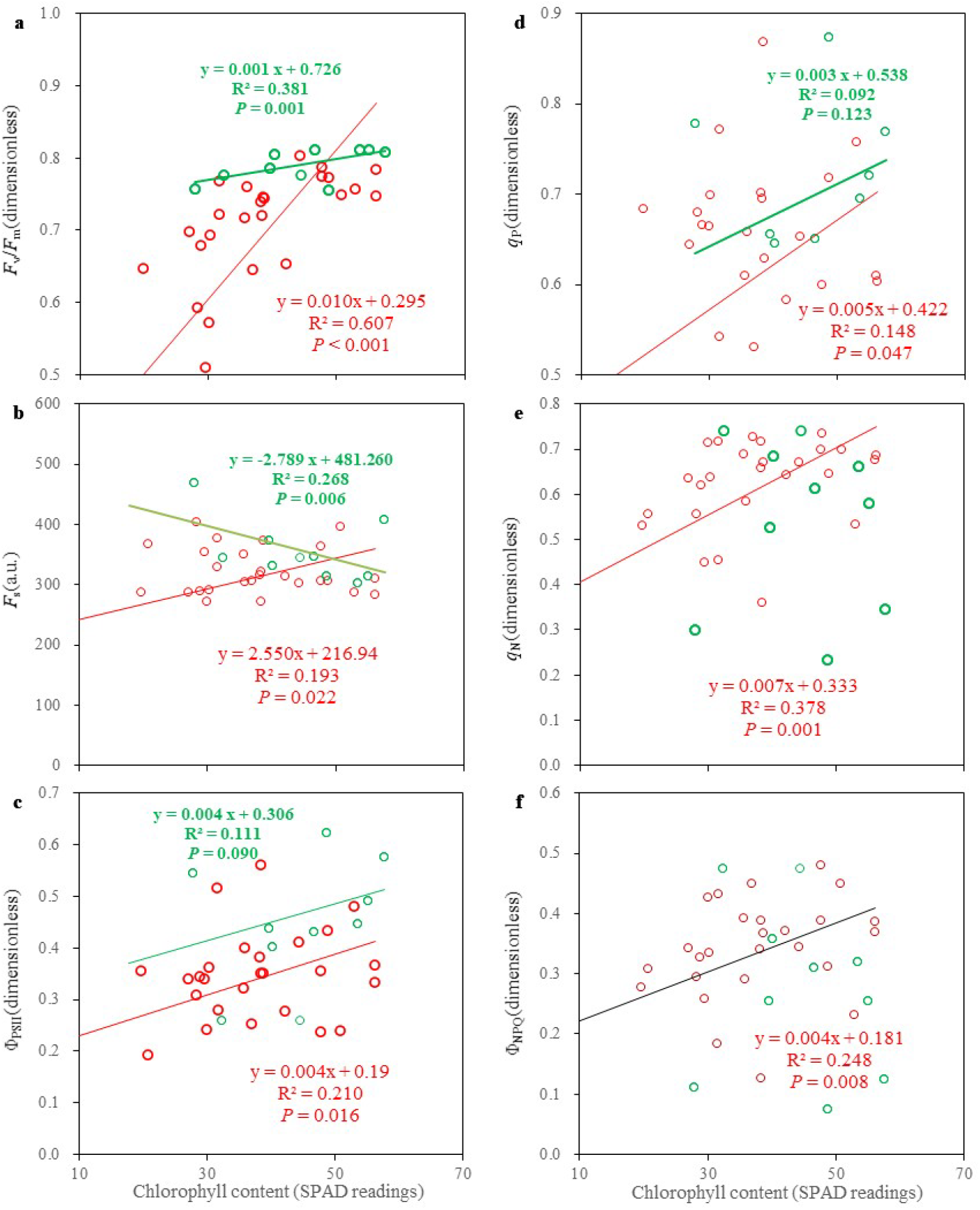
Relationships between fluorescence parameters and chlorophyll content at pre-tasseling (VT, green parts) and post-VT (red parts) stages in maize field (2015). *F_v_/F_m_*, maximal quantum efficiency of photosystem II (PSII); *F*_s_, steady-state fluorescence; Φ_PSII_, the yield of PSII photochemistry; *q*_p_, photochemical quenching based on puddle model; *q*_N_, non-photochemical quenching; Φ_NPQ_, light-induced regulated non-photochemical quenching.

We also performed a PCA to test the relationships between functional traits and the different patterns (Fig. 8). The first two principal components (PCs) accounted for 70.1 % of the total variations. The loadings of SPAD, *F*_v_/*F*_m_, *F*_m_, *F_O_* and 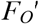 were in quadrant I, while those of Φ_PSII_, 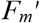 *q*_P_ and *q*_L_ were in quadrant II. The markers most representative of non-photochemical quenching traits (e.g., NPQ, *q*_N_ and Φ_NPQ_) in relation to non-photochemical radiative energy dissipation capability were sorted into quadrant III. Projection on the treatment effects showed that the three irrigation regimes were sorted in the coordinate plane, with control treatment mostly in quadrant II, and SD scattered in all four quadrants (Fig. 8).

**Fig. 8.**
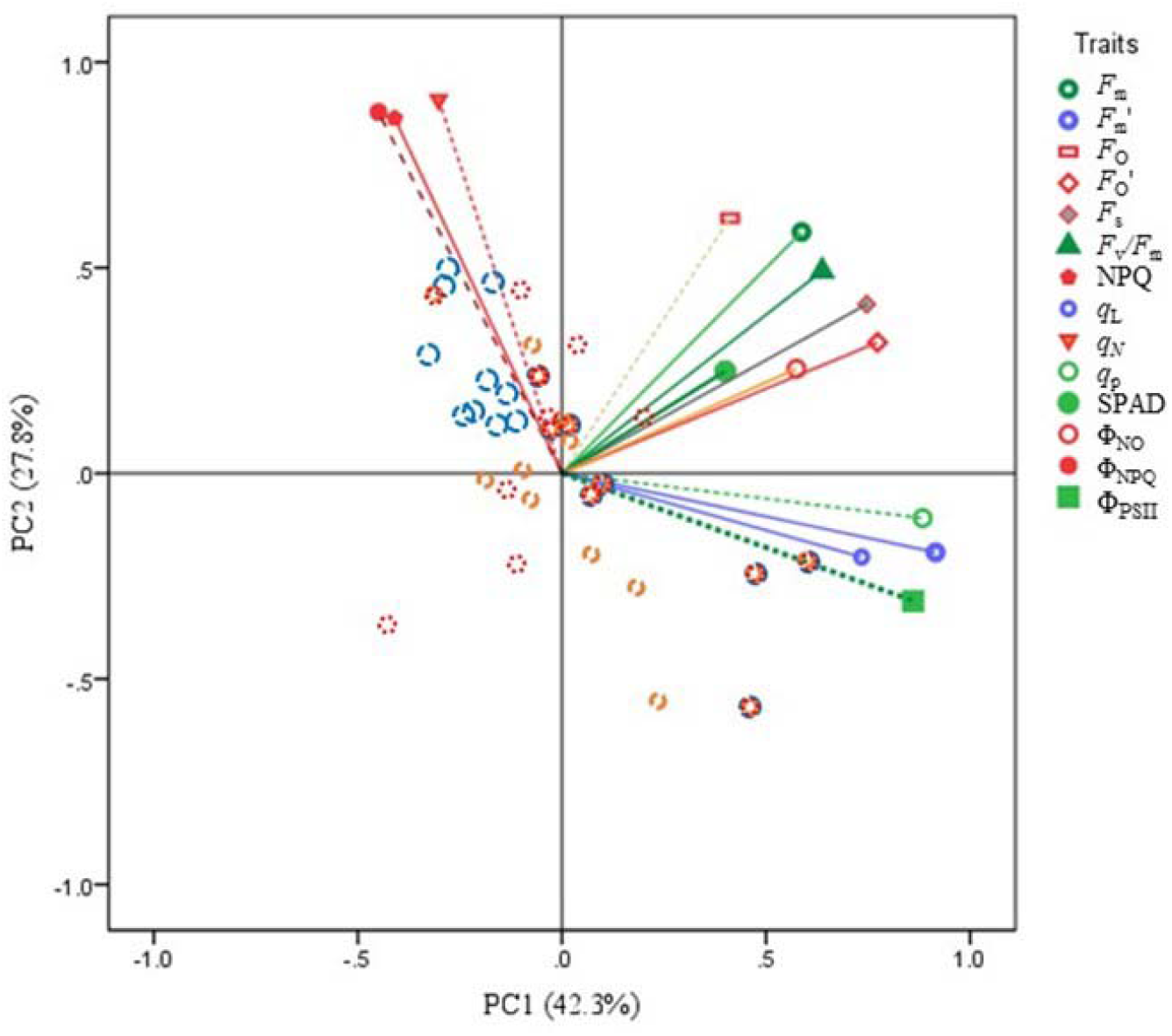
Principal component analysis on plant functional traits under the three irrigation regimes [i.e., control, moderate drought (MD), and severe drought (SD)]. The traits’ loadings on the first two principal components (PCs) are shown, and their projections are sorted by the three irrigation regimes. Dotted green, orange, and red circles represent the PC scores of control, MD, and SD treatments, respectively. *F_m_*, maximal fluorescence yield; 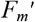, maximal light-adapted fluorescence yield; *F_o_*, minimal fluorescence yield; 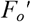, minimal fluorescence at light-adapted state; *F*_s_, steady-state fluorescence; *F_v_F_m_*, maximal quantum efficiency of photosystem II (PSII);Φ_PSII_, yield of PSII photochemistry; *q*_p_, photochemical quenching based on puddle model; *q*_L_, photochemical quenching based on lake models, *q*_N_, non-photochemical quenching; Φ_NPQ_, light-induced regulated non-photochemical quenching; Φ_NO_, quantum yield of non-regulated energy loss.

## Discussion

Water cycle changes could substantially impact plant growth, photosynthetic processes and many crucial physiological functions and nutrient status, thereby affecting plant productivity and crop yield (e.g., Izanloo *et al.*, 2008; Lobell *et al.*, 2014; Kimm *et al.*, 2020). Drought and rewetting may often occur at intervals and are predicted to happen more frequently and severely under climatic change (Dai, 2012; IPCC, 2014; Donat *et al.*, 2016; Diffenbaugh *et al.*, 2017). Indeed, sporadic precipitation is a critical issue in maintaining ecosystem productivity and its structural stability, particularly in arid and semi-arid areas and/or in rain-fed planting regions (Reynolds, 2004; Cooper *et al.*, 2008; Song *et al.*, 2018; Guo and Ogle, 2019).

Maize plays a critical role in meeting the global food demands and is one of the most widely planted staple crops worldwide (Haarhoff and Swanepoel, 2018; FAO, 2020). In this study, the greenhouse and field experiments demonstrated how maize photosynthetic functional traits respond to the abnormal precipitation alterations, including drought, flooding, and rewatering at different growth stages, thereby providing key information for managing crop production. Our main findings were that 1) drought and flooding severely hampered photosynthetic capacity in maize plants, particularly under severe and/or long water stress, in support of the first hypothesis; 2) rewatering could result in partial recovery of some photosynthetic traits, with few compensatory responses, in partial support of our second hypothesis; and 3) the photosynthetic resilience to drought was dependent on the drought severity and the plant developmental stage, largely supporting the third hypothesis. These findings can shed light on ways to improve regulation of crop functional traits to deal with erratic precipitation regimes and may lead to better breeding and management practices for crops that have high drought resistance and drought-resilience traits (Kromdijk *et al.*, 2016; Song *et al.*, 2018; Gupta *et al.*, 2020).

### Drought and flooding

In agreement with previous work (e.g., Chaves *et al.*, 2003, 2009; Xu *et al.*, 2006; Xu and Zhou, 2009; Gupta *et al.*, 2020), the results of the present study indicated that severe drought stress can substantially reduce photosynthetic capacity, as characterised by declines in chlorophyll content, net light-saturated photosynthetic rate (*A*_sat_), stomatal conductance (*g*_s_), and quantum yield of PSII electron transport (Φ_PSII_) in both the greenhouse and field experiments. However, the photosynthetic capacity attenuated more substantially and steeply as the drought stress persisted in our experiment, indicating a strong dependence on the duration, severity and timing of droughts. Thus, only mild or moderate or short drought stresses were conducive to the development of a regulative response of plants for resistance to water deficit in terms of the changes in the root system (e.g., Trapeznikov *et al.*, 2003), leaf thickness (Sack and Grubb, 2002) and antioxidative defence system (Foyer and Noctor, 2005). This observation may aid in implementations of deficit irrigation, water saving agriculture, and sustainable development (Fereres and Soriano, 2007; Geerts and Raes, 2009; Du *et al.*, 2015; Silveira *et al.*, 2020; Kimm *et al.*, 2020).

The present findings also demonstrated that flooding led to a decline in SPAD and *A*_sat_, but not *g*_s_. Indeed, under an anaerobic environment, plants may have adaptive responses to flooding stress that include aerenchyma formation in the roots and the development of adventitious roots (Mano *et al.*, 2006), alteration of the profile of protein synthesis related to anoxic tolerance (Subbaiah and Sachs, 2003), and enhanced starch accumulation (Mutava *et al.*, 2015). An involvement of ethylene regulation is also associated with an enhancement of photochemical and non-photochemical radiative energy dissipation capability (De Pedro *et al.*, 2020). Our results also indicated a higher tolerance of maize to flooding stress in terms of Φ_PSII_ and photochemical quenching (*q*_P_), relative to drought stress, highlighting the distinct effects of these two stresses (Mutava *et al.*, 2015; Zhu *et al.*, 2020). An antagonistic effect on *g*_s_ has been reported (see also Zhu *et al.*, 2020). Maintaining stomatal opening may promote water release to alleviate the stress due to excessive water, again highlighting the positive regulation in response to anoxic conditions (Zhu *et al.*, 2020).

### Recovery and resilience

As previously reported, a depression in photosynthesis potentials by a previous drought can be markedly stimulated by rewetting; however, whether or how much these potentials recover depends on drought intensity and/or the persistence period (e.g., Xu *et al.*, 2009, 2010; Creek *et al.*, 2018). In the current experiment, partial, full and over recovery of photosynthetic traits were all observed in terms of both recovery and resilience indices, specifically depending on the duration and persistence of the drought, the plant developmental stages and the different functional traits, as well as the crop species and cultivar (Figs 1-6; Xu *et al.*, 2009; Creek *et al.*, 2018). For instance, an over-compensatory recovery (i.e., a positive percentage of the resilience) in *g*_s_ was observed in maize (Figs. 2b, 3b); however, *g*_s_ only achieved a partial recovery in a grass species (Xu *et al.*, 2009). Creek e*t al*. (2008) recently reported that, after rewatering, the *A*_net_ of a semiarid species can return to the pre-drought stress level within 2-4 weeks, whereas *g*_s_ performs a slower recovery. A recent report by Johnson *et al.* (2018) indicated that photosynthesis was not fully recovered in wheat plants because of the photosynthetic damage due to hydraulic decline in the leaves subjected to drought. Increased embolism is tightly related to a complete lack of photosynthetic recovery. However, Creek *et al.* (2018) found that photosynthetic recovery can be decoupled from the recovery of plant hydraulics, indicating that the impaired hydraulic function throughout the recovery period perhaps does not influence the complete recovery of *A*_net_ from drought. Thus, the underlying mechanism needs to be investigated further.

The enhancement of plant growth due to rewatering has been addressed by many researchers (Reynolds *et al.*, 2004; Siopongco *et al.*, 2006; Xu *et al.*, 2009; Song *et al.*, 2018). As recently reported by Abid *et al.* (2018), tolerant wheat plants showing high photosynthetic capacity during drought and rapid recovery after re-irrigating did not show marked yield declines relative to the sensitive cultivars, indicating that the plant’s ability to maintain/restore growth and physiological functions during pre/post-drought in the vegetative period might play a crucial role in determining crop productivity. Upon rewatering, the rapid growth of new tissues, such as a new leaf, might accelerate plant growth, potentially enhancing CO_2_ assimilation (Pinheriro *et al.*, 2004). This may be a result of positive source–sink interactions, as a strong sink requirement (e.g. new tissue) can enhance the carbon assimilation rate (Minchin and Lacointe, 2005; White *et al.*, 2015; Parvin *et al.*, 2020). Higher resilience of *A*_sat_ and *g*_s_ occurred in the younger leaves relative to mature ones, implying a greater ability to recover in the new leaves that serve as both active source–sink organs (Figs 2, 3; Roitsch, 1999). The maize plants were exposed to drought stress for only several days, so leaf length after rewatering was restored to a similar level to that of the control plants, indicating no occurrence of overcompensation (Acevedo *et al.*, 1971; Xu *et al.*, 2009; Hofer *et al.*, 2017). Thus, the extent of compensation for drought by the triggering of new tissues following rewatering might determine the final plant/crop production and would depend strongly on the severity, duration, and timing of the drought stress (Hsiao, 1973; Xu *et al.*, 2009; Hofer *et al.*, 2017).

### Associations between functional traits

The distinct responses of the functional traits such as *A*_sat_ and *g*_s_ to drought, flooding and rewetting suggested that coordinated associations between the functional traits could reveal the underlying mechanism (see also Creek *et al.*, 2018). For instance, the SPAD reading (e.g., Ciganda *et al.*, 2009), as an indicator of relative chlorophyll concentration, has different associations with photosynthetic function activities at different plant development stages: stronger associations were observed post-VT (tasselling stage) than pre-VT (Fig. 7). This might indicate that a greater coupling relationship appears at later plant developmental stages and that leaf age/senescence could be involved in the responses to drought and rewatering. This finding may further improve our understanding of how plants respond to water status changes at different developmental stages. For instance, many previous studies have indicated that drought damage increases, while tolerance decreases, with increasing senescence (e.g., David *et al.*, 1998; Shah and Paulsen, 2003; Chaves *et al.*, 2003; Xu *et al.*, 2008; Jiang *et al.*, 2020). However, rewatering may lessen the senescence processes (Xu *et al.*, 2010; Jiang *et al.*, 2020), thereby leading to changes in associations between functional traits such as coupling and trade-off occurrences. Moreover, as revealed by the PCA (Fig.8), a distinct pattern of loadings between Φ_PSII_, Φ_NPQ_ and Φ_NO_ highlights a feasible trade-off strategy by balancing the yields among photochemical processes for the energy absorbed by PSII, dissipation of non-photochemical responses and other non-photochemical losses, which would involve the xanthophyll cycle and PsbS protein expression (Murchie and Lawson, 2013; Kromdijk *et al.*, 2016; Sacharz *et al.*, 2017).

## Conclusions

Quantifying and defining plant functional traits to assess and predict drought effects and post-drought recovery are relevant issues due to the pressing needs imposed by climate change (e.g., Creek *et al.*, 2018; Gupta *et al.*, 2020). In this study, we conducted greenhouse and field experiments to explore how maize photosynthetic functional traits respond to drought, flooding, and rewatering at different growth stages. The main findings were the following: 1) drought and flooding constrains photosynthetic capacity, particularly under severe and/or long water stress; 2) rewatering results in a partial recovery of most of the photosynthetic traits, with few compensatory responses; 3) photosynthetic resilience to drought following rewetting strongly depends on the drought severity and its persistence and duration, as well as the plant growth stage. The distinct responses of various functional traits to drought, flooding, and rewetting can translate to a regulative strategy of trade-off. The coordinated changes in chlorophyll content, gas exchange, fluorescence parameters (quantum efficiency of PSII, and photochemical and non-photochemical radiative energy dissipation) may largely contribute to the enhancements of drought resistance and resilience of plants. The associations of chlorophyll concentrations with photosynthetic functional activities were also stronger post-VT than pre-VT, implying that leaf age/senescence may be involved in the responses to drought and rewatering. These findings may further improve our understanding of how plants respond to water status changes at different plant developmental stages. This knowledge can be helpful for breeding crops with high drought-resistant and drought-resilience traits and for establishing management practices when facing climate change (e.g., Kromdijk *et al.*, 2016; Leakey *et al.*, 2019; Kimm *et al.*, 2020; Gupta *et al.*, 2020).

## Acknowledgements

This study was jointly funded by the National Key Research and Development Program of China (2016YFD0300106), National Natural Science Foundation of China (31661143028), and China Special Fund for Meteorological Research in the Public Interest (GYHY201506001-3). The authors are grateful to Yuhui Wang, Bingrui Jia, Yanling Jiang, and Jian Song for their help during the study. The authors are also grateful to Hongying Yu, Quanhui Ma, and Liang Li for their loyal assistances during the two experiments.

## Author contributions

Zhenzhu Xu and Guagnsheng Zhou deceived and designed the study; Miao Qi, Xiaodi Liu, Yibo Li, He Song, and Feng Zhang conducted the experiment works. Miao Qi, Xiaodi Liu, Yibo Li, and Zhenzhu Xu performed the data analyses. All authors wrote and proofread the manuscript.

## Availability of data and materials

The data sets supporting the results of this article are included within the article and its supporting information file.

## References

Abid M, Ali S, Qi LK, Zahoor R, Tian Z, Jiang D, Snider JL, Dai T. 2018. Physiological and biochemical changes during drought and recovery periods at tillering and jointing stages in wheat *(Triticum aestivum* L.). Scientific Reports 8, 4615.

Acevedo E, Hsiao TC, Henderson DW. 1971. Immediate and subsequent growth responses of maize leaves to changes in water statues. Plant Physiology 48, 631–636.

Ben-Ari T, Adrian J, Klein T, Calanca P, Van der Velde M, Makowski D. 2016. Identifying indicators for extreme wheat and maize yield losses. Agricultural and Forest Meteorology 220, 130–140.

Bhaskar R, Arreola F, Mora F, Martinez-Yrizar A, Martinez-Ramos M, Balvanera P. 2018. Response diversity and resilience to extreme events in tropical dry secondary forests. Forest Ecology and Management 426, 61–71.

Chaves MM, Flexas J, Pinheiro C. 2009. Photosynthesis under drought and salt stress: regulation mechanisms from whole plant to cell. Annal of Botany 103, 551–560.

Chaves MM, Maroco JP, Pereira JS. 2003. Understanding plant responses to drought—from genes to the whole plant. Functional Plant Biology 30, 239–264.

Ciganda V, Gitelson A, Schepers J. 2009. Non-destructive determination of maize leaf and canopy chlorophyll content. Journal of Plant Physiology 166, 157–167.

Cooper PJM, Dimes J, Rao KPC, Shapiro B, Shiferaw B, Twomlow S. 2008. Coping better with current climatic variability in the rain-fed farming systems of sub-Saharan Africa: An essential first step in adapting to future climate change? Agriculture, Ecosystems & Environment 126, 24–35.

Creek D, Blackman C, Brodribb TJ, Choat B, Tissue DT. 2018. Coordination between leaf, stem and root hydraulics and gas exchange in three arid-zone angiosperms during severe drought and recovery. Plant, Cell and Environment 41, 2869–2881.

Dai A. 2012. Increasing drought under global warming in observations and models. Nature Climate Change 3, 52–58.

Daryanto S, Wang L, Jacinthe PA. 2016. Global synthesis of drought effects on maize and wheat production. PloS one 11, 5.

David MM, Coelho D, Barrote I, Correia MJ. 1998. Leaf age effects on photosynthetic activity and sugar accumulation in droughted and rewatered *Lupinus albus* plants. Australian Journal of Plant Physiology 25, 299–306.

De Pedro LF, Mignolli F, Scartazza A, Melana Colavita JP, Bouzo CA, Vidoz ML. 2020. Maintenance of photosynthetic capacity in flooded tomato plants with reduced ethylene sensitivity. Physiologia Plantarum doi:10.1111/ppl.13141.

Diffenbaugh NS, Singh D, Mankin JS, Horton DE, Swain DL, Touma D, Charland A, Liu Y, Haugen M, Tsiang M, Rajaratnam B. 2017. Quantifying the influence of global warming on unprecedented extreme climate events. Proceedings of the National Academy of Sciences 114, 4881–4886.

Du T, Kang S, Zhang J, Davies WJ. 2015. Deficit irrigation and sustainable water-resource strategies in agriculture for China’s food security. Journal of Experimental Botany 66, 2253–2269.

FAO. 2020. FAOSTAT database. FAO, Rome. http://www.fao.org/faostat/en/#data/QC (accessed on 19 June 2020).

Fereres E, Soriano MA. 2007. Deficit irrigation for reducing agricultural water use. Journal of Experimental Botany 58, 147–259.

Foyer CH, Noctor G. 2005. Oxidant and antioxidant signaling in plants: a revaluation the concept of oxidative stress in a physiological context. Plant, Cell and Environment 28, 1056–1071.

Geerts S, Raes D. 2009. Deficit irrigation as an on-farm strategy to maximize crop water productivity in dry areas. Agricultural water management 96, 1275–1284.

Guo JS, Ogle K. 2019. Antecedent soil water content and vapor pressure deficit interactively control water potential in *Larrea tridentate*. New Phytologist 221, 218–232.

Gupta, A., Rico-Medina, A., & Caño-Delgado, A. I. (2020). The physiology of plant responses to drought. Science, 368(6488), 266–269.

Haarhoff SJ, Swanepoel PA. 2018. Plant population and maize grain yield: a global systematic review of rainfed trials. Crop Science 58, 1–11.

Harrison SP, LaForgia ML, Latimer AM. 2018. Climate□driven diversity change in annual grasslands: Drought plus deluge does not equal normal. Global Change Biology 24, 1782–1792.

Hofer D, Suter M, Buchmann N, Lüscher A. 2017. Nitrogen status of functionally different forage species explains resistance to severe drought and post-drought overcompensation. Agriculture, Ecosystems and Environment 236, 312–322.

Holling CS. 1973. Resilience and stability of ecological systems. Annual Review of Ecology and Systematics 4, 1–23.

Hsiao TC. 1973. Plant responses to water stress. Annual Review of Plant Physiology 24, 519–570.

IPCC. 2014. Climate Change 2014: Synthesis Report. Contribution of Working Groups I, II and III to the Fifth Assessment Report of the Intergovernmental Panel on Climate Change (Core Writing Team, Pachauri RK, Meyer LA, eds.). IPCC, Geneva, Switzerland, 151 pp.

Izanloo A, Condon AG, Langridge P, Tester M, Schnurbusch T. 2008. Different mechanisms of adaptation to cyclic water stress in two South Australian bread wheat cultivars. Journal of Experimental Botany 59, 3327–46.

Jiang T, Dou Z, Liu J, Gao Y, Malone RW, Chen S, Feng H, Yu Q, Xue G, He J. 2020. Simulating the influences of soil water stress on leaf expansion and senescence of winter wheat. Agricultural and Forest Meteorology 291, 108061.

Johnson KM, Jordan GJ, Brodribb TJ. 2018. Wheat leaves embolised by water stress do not recover function upon rewatering. Plant, Cell and Environment 41, 2704–2714.

Kimm H, Guan K, Gentine P, Wu J, Bernacchi CJ, Sulman BN, Griffis TJ, Lin C. 2020. Redefining droughts for the US Corn Belt: The dominant role of atmospheric vapor pressure deficit over soil moisture in regulating stomatal behavior of Maize and Soybean. Agricultural and Forest Meteorology 287, 107930.

Kramer DM, Johnson G, Kiirats O, Edwards GE. 2004. New fluorescence parameters for the determination of QA redox state and excitation energy fluxes. Photosynthesis Research 79, 209–218.

Kromdijk J, Głowacka K, Leonelli L, Gabilly ST, Iwai M, Niyogi KK, Long SP. 2016. Improving photosynthesis and crop productivity by accelerating recovery from photoprotection. Science 354, 857–861.

Leakey AD, Ferguson JN, Pignon CP, Wu A, Jin Z, Hammer GL, Lobell DB. 2019. Water use efficiency as a constraint and target for improving the resilience and productivity of C_3_ and C_4_ crops. Annual Review of Plant Biology 70, 781–808.

Li Y, Song H, Zhou L, Xu Z, Zhou G. 2019. Vertical distributions of chlorophyll and nitrogen and their associations with photosynthesis under drought and rewatering regimes in a maize field. Agricultural and Forest Meteorology 272, 40–54.

Li Z, Sun Z. 2016. Optimized single irrigation can achieve high corn yield and water use efficiency in the Corn Belt of Northeast China. European Journal of Agronomy, 75, 12–24.

Liu Z, Yang X, Hubbard KG, Lin X. 2012. Maize potential yields and yield gaps in the changing climate of northeast China. Global Change Biology 18, 3441–3454.

Lobell DB, Roberts MJ, Schlenker W, Braun N, Little BB, Rejesus RM, Hammer GL. 2014. Greater sensitivity to drought accompanies maize yield increase in the US Midwest. Science 344, 516–519.

Mano Y, Omori F, Takamizo T, Kindiger B, Bird RM, Loaisiga CH. 2006. Variation for root aerenchyma formation in flooded and non-flooded maize and teosinte seedlings. Plant and Soil 281, 269–279.

Maxwell K, Johnson GN. 2000. Chlorophyll fluorescence—a practical guide. Journal of Experimental Botany 51, 659–668.

Müller F, Bergmann M, Dannowski R, Dippner JW, Gnauck A, Haase P, Jochimsen M, Kasprzak P., Kröncke I, Kümmerlin R, Küster M. 2016. Assessing resilience in long-term ecological data sets. Ecological Indicators 65, 10–43.

Mutava RN, Prince SJK, Syed NH, Song L, Valliyodan B, Chen W, Nguyen HT. 2015. Understanding abiotic stress tolerance mechanisms in soybean: a comparative evaluation of soybean response to drought and flooding stress. Plant Physiology and Biochemistry 86, 109–120.

Myers, SS, Smith, MR, Guth, S., Golden, CD, Vaitla, B., Mueller, ND, Dangour AD, Huybers P. 2017. Climate change and global food systems: potential impacts on food security and undernutrition. Annual Review of Public Health, 38, 259–277.

Parvin S, Uddin S, Tausz-Posch S, Armstrong R, Tausz M. 2020. Carbon sink strength of nodules but not other organs modulates photosynthesis of faba bean *(Vicia faba)* grown under elevated [CO_2_] and different water supply. New Phytologist 227, 132–145.

Pinheriro C, Passarinho JA, Ricardo CP. 2004. Effect of drought and rewatering on the metabolism of *Lupinus albus* organs. Journal of Plant Physiology 161, 1203–1210.

Resilience Alliance. 2020. http://www.resalliance.org/about.

Reynolds JF, Kemp PR, Ogle K, Fernández RJ. 2004. Modifying the ‘pulse-reserve’ paradigm for deserts of North America: precipitation pulses, soil water and plant responses. Oecologia 141, 194–210.

Roitsch T. 1999. Source-sink regulation by sugar and stress. Current Opinion in Plant Biology 2, 198–206.

Rosa L, Chiarelli DD, Rulli MC, Dell’Angelo J, D’Odorico P. 2020. Global agricultural economic water scarcity. Science Advances 6, eaaz6031.

Roudier P, Andersson JC, Donnelly C, Feyen L, Greuell W, Ludwig F. 2016. Projections of future floods and hydrological droughts in Europe under a +2°C global warming. Climatic Change 135, 341–355.

Ruppert JC, Harmoney K, Henkin Z, Snyman HA, Sternberg M, Willms W, Linstädter A. 2015. Quantifying drylands’ drought resistance and recovery: the importance of drought intensity, dominant life history and grazing regime. Global Change Biology 21, 1258–1270.

Sacharz J, Giovagnetti V, Ungerer P, Mastroianni G, Ruban A. 2017. The xanthophyll cycle affects reversible interactions between PsbS and light-harvesting complex II to control non-photochemical quenching. Nature Plants 3, 16225.

Sack L, Grubb PJ. 2002 The combined impacts of deep shade and drought on the growth and biomass allocation of shade-tolerant woody seedlings. Oecologia 131, 175–185.

Schreiber UBWN, Bilger W, Neubauer C. 1994. Chlorophyll fluorescence as a nonintrusive indicator for rapid assessment of in vivo photosynthesis. Ecophysiology of Photosynthesis 100, 49–70.

Shah NH, Paulsen GM. 2003. Interaction of drought and high temperature on photosynthesis and grain-filling of wheat. Plant and Soil 257, 219–226.

Sharp RE, Poroyko V, Hejlek LG, Spollen WG, Springer GK, Bohnert HJ, Nguyen HT. 2004. Root growth maintenance during water deficits: physiology to functional genomics. Journal of Experimental Botany 55, 2343–2351.

Silveira LK, Pavão GC, dos Santos Dias CT, Quaggio JA, de Matos Pires RC. 2020. Deficit irrigation effect on fruit yield, quality and water use efficiency: A long-term study on Pêra-IAC sweet orange. Agricultural Water Management 231, 106019.

Siopongco JDLC, Yamauchi A, Salekdeh H, Bennett J, Wade LJ. 2006. Growth and water use response of doubled-haploid rice lines to drought and rewatering during the vegetative stage. Plant Production Science 9, 141–151.

Song H, Li Y, Zhou L, Xu Z, Zhou G. 2018. Maize leaf functional responses to drought episode and rewatering. Agricultural and Forest Meteorology 249, 57–70.

Subbaiah CC, Sachs MM. 2003. Molecular and cellular adaptations of maize to flooding stress. Annals of Botany 91, 119–127.

Sun CX, Li CC, Zhang CY, Hao LY, Song M, Liu W, Zhang YL. 2018. Reflectance and biochemical responses of maize plants to drought and re-watering cycles. Annals of Applied Biology 172, 332–345.

Trapeznikov VK, Ivanov II, Kudoyarova GR. 2003. Effect of heterogeneous distribution of nutrients on root growth, ABA content and drought resistance of wheat plants. Plant and Soil, 252, 207–214.

Trenberth KE, Dai A, Van Der Schrier G, Jones PD, Barichivich J, Briffa KR, Sheffield J. 2014. Global warming and changes in drought. Nature Climate Change 4, 17–22.

Van Ruijven J, Berendse, F. 2010. Diversity enhances community recovery, but not resistance, after drought. Journal of Ecology 98, 81–86.

Voronin PY, Maevskaya SN, Nikolaeva MK. 2019. Physiological and molecular responses of maize (*Zea mays* L.) plants to drought and rehydration. Photosynthetica 57, 850–856.

White AC, Rogers A, Rees M, Osborne CP. 2015. How can we make plants grow faster? A source-sink perspective on growth rate. J Exp Bot 67, 31–45. https://doi.org/10.1093/jxb/erv447.

Xu Z, Shimizu H, Ito S, Yagasaki Y, Zou C, Zhou G, Zheng Y. 2014. Effects of elevated CO_2_, warming and precipitation change on plant growth, photosynthesis and peroxidation in dominant species from North China grassland. Planta 239, 421–435.

Xu Z, Zhou G, Shimizu H. 2009. Are plant growth and photosynthesis limited by pre-drought following rewatering in grass? Journal of Experimental Botany 60, 3737–3749.

Xu Z, Zhou G. 2011. Responses of photosynthetic capacity to soil moisture gradient in perennial rhizome grass and perennial bunchgrass. BMC Plant Biology 11, 21.

Xu ZZ, Zhou GS, Shimizu H. 2009. Are plant growth and photosynthesis limited by pre-drought following rewatering in grass? Journal of Experimental Botany 60:3737–3749.

Xu ZZ, Zhou GS, Shimizu H. 2010. Plant responses to drought and rewatering. Plant Signaling & Behavior 5: 649–654.

Xu ZZ, Zhou GS, Wang YL, Han GX, Li YJ. 2008. Changes in chlorophyll fluorescence in maize plants with imposed rapid dehydration at different leaf ages. Journal of Plant Growth Regulation 27, 83–92.

Xu ZZ, Zhou GS. 2007. Photosynthetic recovery of a perennial grass *Leymus chinensis* after different periods of soil drought. Plant Production Science 10, 277–285.

Zhu R, Wu F, Zhou S, Hu T, Huang J, Gao Y. 2020. Cumulative effects of drought–flood abrupt alternation on the photosynthetic characteristics of rice. Environmental and Experimental Botany 169, 103901.

